# Cognitive behavioral phenotyping of *DSCAM* heterozygosity as a model for autism spectrum disorder

**DOI:** 10.1101/2024.06.03.597158

**Authors:** Ryan C. Neff, Katherine A. Stangis, Ujjawal Beniwal, Ty Hergenreder, Bing Ye, Geoffrey G. Murphy

## Abstract

It is estimated that 1 in 36 children are affected by autism spectrum disorder (ASD) in the United States, which is nearly a twofold increase from a decade ago. Recent genetic studies have identified *de novo* loss-of-function (dnLoF) mutations in the *Down Syndrome Cell Adhesion Molecule (DSCAM)* as a strong risk factor for ASD. Previous research has shown that *DSCAM* ablation confers social interaction deficits and perseverative behaviors in mouse models. However, it remains unknown to what extent *DSCAM* underexpression captures the full range of behaviors, specifically cognitive phenotypes, presented in ASD. Here, we conducted a comprehensive cognitive behavioral phenotyping which revealed that loss of one copy of *DSCAM*, as in the *DSCAM*^2J^+/− mice, displayed hyperactivity, increased anxiety, and motor coordination impairments. Additionally, hippocampal-dependent learning and memory was affected, including working memory, long-term memory, and contextual fear learning. Interestingly, implicit learning processes remained intact. Therefore, *DSCAM* LoF produces autistic-like behaviors that are similar to human cases of ASD. These findings further support a role for *DSCAM* dnLoF mutations in ASD and suggest *DSCAM*^2J^+/− as a suitable model for ASD research.

**Summary Statement:** Autism spectrum disorder represents a growing patient population. Loss of one copy of the *DSCAM* gene provides a promising mouse model that reproduces autistic-like behaviors for research and therapeutic testing.

## Introduction

Over the last decade, autism spectrum disorder (ASD) has been increasing in prevalence considerably, and it is now estimated that 1 in 36 children born in the United States meet the diagnostic criteria for the disorder (Maenner et al., 2023). Autism spectrum disorder is characterized by an array of symptoms including somatosensory abnormalities as well as deficits in several social and cognitive domains. Specifically, children and adults diagnosed with ASD can exhibit hyperactivity and heightened anxiety (Avni et al., 2018); motor coordination difficulties (Miller et al., 2014); social and cognitive impairments, including developmental delays (Huguet et al., 2013, Charman et al., 2011, Shan et al., 2022); and fear learning deficits (Powell et al., 2016). These symptoms often manifest on a spectrum, as the name implies, and therefore affect individuals with differing severity. Furthermore, the causes of most ASD cases are currently unknown. While research had historically focused on environmental circumstances contributing to ASD onset and exacerbation (Bölte et al., 2019), the current focus of the field has shifted to genetic risk factors (Rylaarsdam and Guemez-Gamboa, 2019, Hodges et al., 2020). Indeed, ASD has been found to accumulate within families and appears to be highly heritable (Sandin et al., 2017, Tick et al., 2016). Genome-wide association studies (GWAS) have found numerous genes to be linked to ASD as risk factors, many of which reside in pathways associated with neurodevelopment and synaptic transmission (Krishnan et al., 2016, Moyses-Oliveira et al., 2020). One gene that has been repeatedly identified as a significant risk factor of ASD is the *Down Syndrome Cell Adhesion Molecule* (*DSCAM*) (Stephan et al., 2015, Stessman et al., 2017, Tychele et al., 2016, Wang et al., 2016, Iossifov et al., 2014). Many of the *DSCAM* mutations that have been discovered are *de novo* loss-of-function (dnLoF) mutations (Iossifov et al., 2014), which likely result in heterozygous *DSCAM* expression. These findings suggest *DSCAM* dnLoF mutation as a potential cause of ASD in the general population.

First characterized in *Drosophila*, *Dscam* is a transmembrane protein that mediates self-avoidance of the neurites of a neurons (Matthews et al., 2007, Soba et al., 2007, Hughes et al., 2007), regulates axonal guidance (Schmucker et al., 2000, Andrews et al., 2008), and promotes the growth of presynaptic terminals (Kim et al., 2013). In the mouse retina, *DSCAM* determines spatial arrangement of neurons during initial growth and subsequent refinement of projections (Fuerst et al., 2008, Li et al., 2015, Fuerst et al., 2010). Moreover, *DSCAM* controls the axon growth of retinal ganglion cells (Bruce et al., 2017). Interestingly, the function of *Dscam*/*DSCAM* appears to act in a dose-dependent manner, as seen in presynaptic growth in both *Drosophila* and mice (Kim et al., 2013, Liu et al., 2023). This has led *DSCAM* to be studied in the context of multiple neuropsychiatric disorders (Stachowicz, 2018). For instance, overexpression of *DSCAM*, which occurs in Down syndrome, has been shown to drive an increase in inhibitory connections in the cortex of Down syndrome mouse models (Liu et al., 2023). On the other hand, a decrease in *DSCAM* expression may underlie some of the key phenotypes that are present in ASD. Dysregulation of NMDA receptors were detected in neurons derived from human iPSCs from an ASD patient where *DSCAM* was deficient, and proper NMDA receptor behavior was restored following the introduction of wild-type *DSCAM* expression (Lim et al., 2021). Complete knockout of *DSCAM* in neurons and astrocytes in mice accelerates dendritic spine development, which may affect initial synapse formation and maturation during post-natal growth (Chen et al., 2022). Moreover, *DSCAM* knockout mice appear to replicate some of the repetitive behaviors and social impairments that are associated with ASD (Chen et al., 2022), further promoting *DSCAM* LoF mutations as a potential cause of ASD. However, ASD is often caused by heterozygosity of risk loci as opposed to a complete ablation of gene expression (An and Claudianos, 2016, Iossifov et al., 2014). Whether *DSCAM* heterozygous mice exhibit ASD-like behavioral phenotypes, particularly those related to cognitive paradigms, is an important question and is currently unknown.

Here, we aimed to provide a comprehensive behavioral assessment of *DSCAM* heterozygous mice (*DSCAM*^2J^+/−, i.e., heterozygotes) that would determine the value of this mouse model for ASD research. We focused on murine behaviors that correlated with those described in ASD, namely activity and sensorimotor function, working and long-term memory, fear learning, and procedural learning. We found that *DSCAM* heterozygosity due to a LoF mutation was sufficient to drive hyperactivity and notable motor coordination deficits along with minor impairments in spatial learning and memory. Additionally, these mice show a substantial and lasting deficit in contextual fear learning. Our findings support a decrease in *DSCAM* expression as a possible explanation of ASD-related behaviors and provide strong evidence for a disease model that could be useful for ASD research and therapeutic testing.

## Results

In order to conduct a thorough behavioral phenotyping, cohorts were taken through a series of behavioral tasks and tests. Multiple cohorts were used throughout the study and evaluated in different sets of experiments. The composition of each cohort and which tests they were subjected to are summarized in **Table 1**.

**Table 1.** Summary of cohorts and respective behavioral tests. All cohorts were composed of littermates resulting from a mating cross between *DSCAM*^2J^+/− and C3H. Mice were age-matched and entered behavioral testing at 2-5 months of age. Behavioral tasks were organized according to degree of aversiveness, with any task that includes an electrical foot-shock being run as the terminal experiment. Cohorts that participated in multiple experiments were given at least one week rest between successive tasks.

| Cohort # | Experiments | # of Wild-types | # of Heterozygotes |
| --- | --- | --- | --- |
| Cohort 1 | Open field exploration<br>Morris water maze<br>Fear conditioning | 8<br>(4M, 4F) | 16<br>(7M, 9F) |
| Cohort 2 | Y-maze<br>Passive avoidance | 12<br>(7M, 5F) | 12<br>(4M, 8F) |
| Cohort 3 | Touchscreen pretraining<br>Visual discrimination | 17<br>(10M, 7F) | 12<br>(5M, 7F) |
| Cohort 4 | Rotarod<br>Hot plate | 9<br>(4M, 5F) | 7<br>(2M, 5F) |

### Loss of one copy of *DSCAM* heightens the state of anxiety and promotes hyperactivity

Activity levels in a novel environment were assessed in the open-field exploration task. Exploration and activity were determined as the total distance travelled during the 10-minute test period, while anxiety was measured by the willingness to explore and remain in the inner zone of the arena. Representative exploration paths are presented in **Fig 1A**. When compared to the wild-type controls, the heterozygotes showed increased distance travelled during the test period (**Fig. 1B**; t=2.310, df=21, p=0.0312). Additionally, the time spent in the inner zone of the arena was decreased in the heterozygotes (**Fig. 1C**; t=2.667, df=21, p=0.0144) signaling a possible heightened state of anxiety present. However, there was no difference observed between the groups in the number of crossings between the zones (**Fig. 1D**; t=0.7716, df=21, p=0.4489). Altogether, these results suggest that the heterozygotes are hyperactive and equally motivated to explore the novel environment as the wild-type mice, as seen by similar number of zone crossings, but that they do not spend prolonged periods of time in the inner zone due to increased anxiety-like behavior.

**Figure 1.**
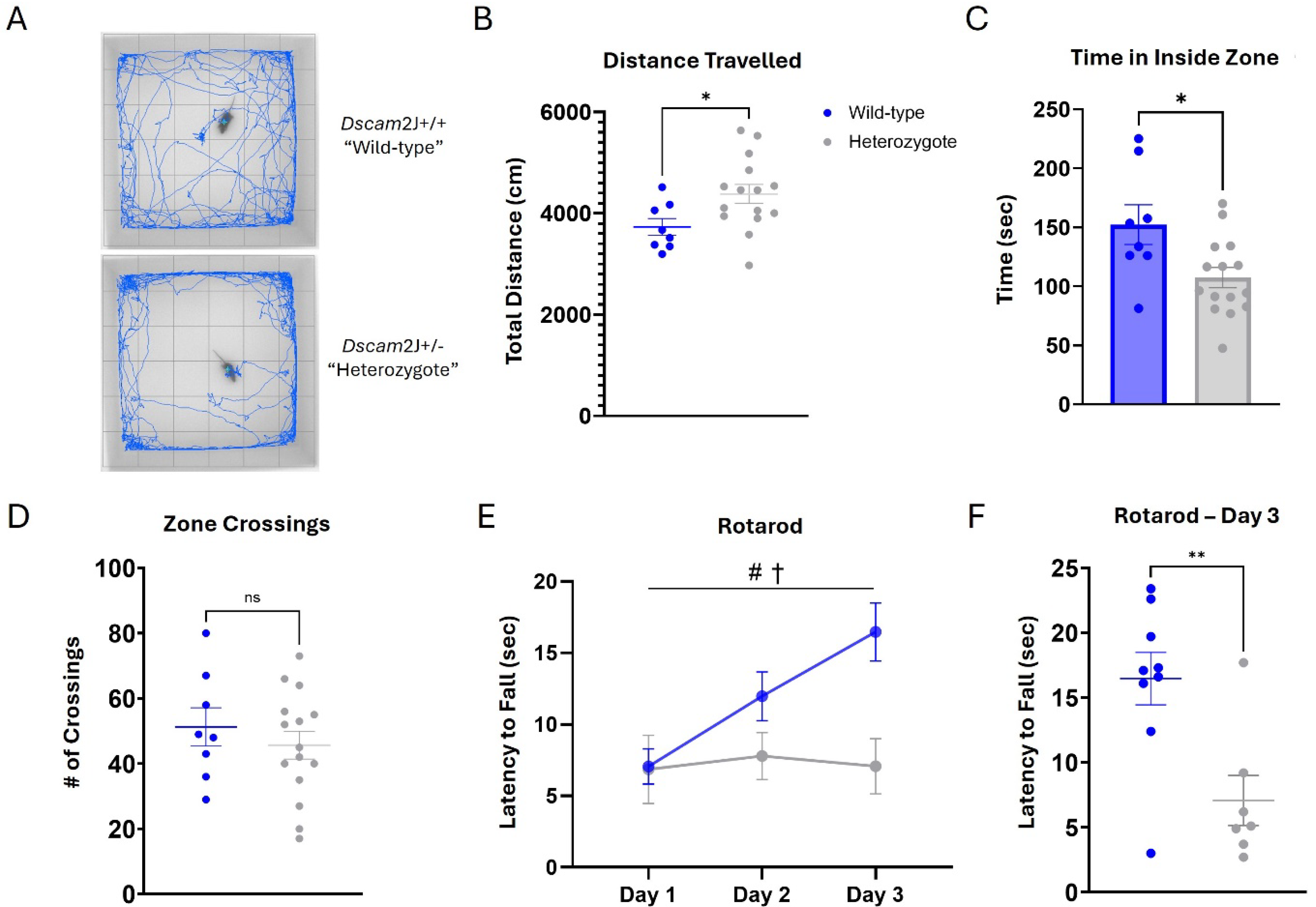
*DSCAM*^2J^+/− heterozygotes exhibit hyperactivity, heightened anxiety, and impaired motor coordination. Exploratory behavior was assessed in the open field during a 10-minute trial. (**A**) Representative trajectories of individual animals demonstrate the discrepancy between genotypes. (**B**) Heterozygotes traveled significantly more distance during exploration. (**C, D**) Heterozygotes spend less time in the inner zone but there was no significant difference in the number of crossings between zones. Locomotor coordination was evaluated on an accelerating rotarod. (**E, F**) Latency to fall was measured, and found to be significantly less in heterozygotes. Results are presented as Mean ± SEM. (**A-D**) Open field *DSCAM*^2J^+/− n = 15 and WT n = 8. Rotarod *DSCAM*^2J^+/− n = 7 and WT n = 9. Statistical tests are as follows: (**B-D, F**) Two-tailed *t*-test. (**E**) Mixed-effects model (REML) where # denotes main effect of time and † denotes main effect of genotype. Significant differences are shown (* p < 0.05 and ** p < 0.01).

### *DSCAM* heterozygotes exhibit impaired locomotor coordination without affecting nociception

As the *DSCAM*^2J^ null has been previously observed to have an altered gait (Fuerst et al., 2010) that could affect their motor performance, locomotor coordination and motor learning was assessed in the *DSCAM*^2J^+/− using the rotarod balancing task. To perform the task, it is necessary that the mice shift and adjust their balance on the rod as it slowly accelerates, and their latency to fall off the rod is recorded. This task was repeated for three consecutive days to allow the mice to learn the task and improve their latency. Unlike the wild-type controls, the heterozygotes showed no improvement across time (**Fig. 1E**; effect of time F (1.552, 20.96) = 5.152, p=0.02; effect of genotype F (1, 14) = 5.214, p=0.039). This was further discovered through an ad-hoc comparison of performance on Day 3 of training, where the heterozygotes had a decreased latency to fall compared to the wild-type (**Fig. 1F**; t=3.279, df=14, p=0.0055), suggesting an impairment in motor coordination.

To examine sensory efficiency, specifically through thermal nociception, the hot plate test was conducted. This test determined the nocifensive behavior response time of each mouse to a heated surface, which was then used as a proxy for the peripheral pain response. There was no difference between the groups in latency to hindlimb lick, indicating that loss of one copy of *DSCAM* produces no overt alterations in thermal nociception (**Fig. S1A**; t=0.4083, df=14, p=0.6892).

### Impaired working memory in *DSCAM* heterozygotes

The Y-maze was used to assess spontaneous alternation as a measure of spatial working memory using visual cues (**Fig. 2A**). As we had previously observed in the open-field exploration task, the heterozygotes displayed hyperactivity in their 8-minute exploration of the Y-maze as well. When compared to the wild-type controls, the heterozygotes had more arm entries (**Fig. 2B**; t=2.249, df=22, p=0.0349) and travelled a greater distance (**Fig. 2C**; t=2.279, df=22, p=0.0327). In alternation rate, the heterozygotes completed fewer correct alternations than chance (50%) would have expected (**Fig. 2D**; wild-type t=0.4863, df=11, p=0.6363; heterozygote t=2.432, df=11, p=0.0333), indicating a greater number of errors and a deficit in spontaneous alternation. Those errors were represented by an increase in direct revisits when compared to wild-type controls (**Fig. 2E**; t=2.433, df=22, p=0.0236), while indirect revisits were similar between the groups (**Fig. 2F**; t=0.9358, df=22, p=0.3595). These findings indicate a significant impairment in the short-term, or working, memory of the *DSCAM* heterozygous mice.

**Figure 2.**
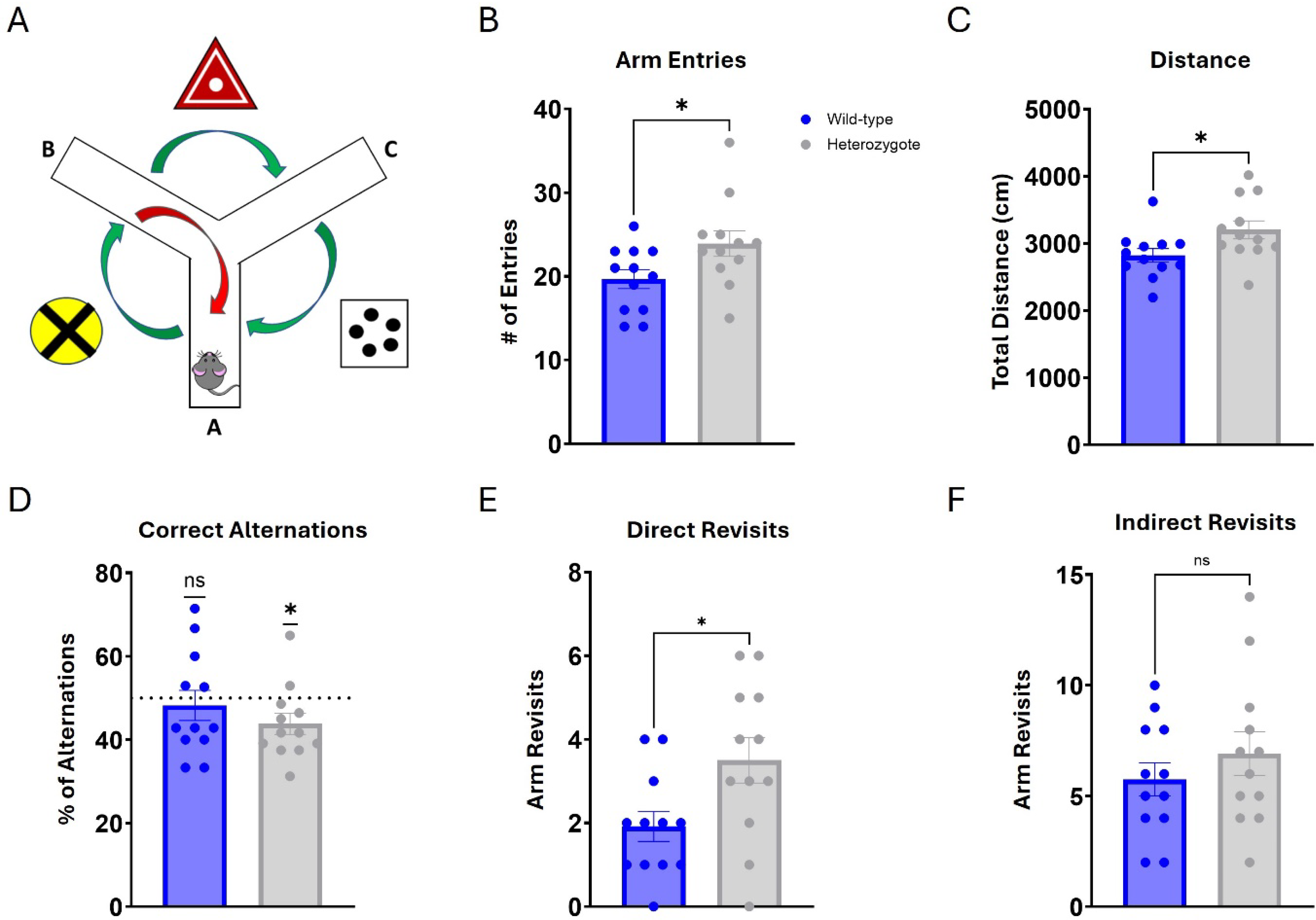
*DSCAM*^2J^+/− heterozygotes are hyperactive and show impairments in spatial working memory. Short-term learning and memory were probed in 8-minute trials in the Y-maze. (**A**) Representation of the maze layout, with three visual cues across from each arm. Correct spontaneous alternations (green arrows) compared to alternation errors (red arrow) were considered as a metric of working memory during maze exploration. (**B, C**) Heterozygotes completed significantly more entries into arms and travelled further during the trial. (**D**) Percentage of triads registered as a correct alternation was significantly below chance (dotted line = 50%) for heterozygotes. (**E, F**) Direct revisit errors were significantly greater in heterozygotes, while indirect revisit errors were shown to be not different between the groups. Results are presented as Mean ± SEM. Sample size: *DSCAM*^2J^+/− n = 12 and WT n = 12. Comparison between heterozygotes and WT mice was analyzed by a two-tailed *t*-test, or a one-tailed *t*-test calculated against chance (50%), and the significant differences are shown (* p < 0.05 and ** p < 0.01).

### Loss of one copy of *DSCAM* impairs memory recall in the Morris water maze task

The Morris water maze tests spatial learning and memory based on environmental cues. The 10-day protocol consists of 9 training days, 3 inter-training memory probes, and visible trials on Day 10 (**Fig. 3A**). Both groups learned the task and improved their training performance across time, with no difference between groups (**Fig. 3B**; effect of time F (5.680, 125.0) = 7.829, p<0.0001; effect of genotype F (1, 22) = 0.1018, p=0.7527). Similarly, both groups decreased their cumulative proximity across training days, to the same extent (**Fig. 3C**; effect of time F (8, 198) = 9.188, p<0.0001; effect of genotype F (1, 198) = 1.339, p=0.2486). Heterozygotes showed a significant increase in distance travelled during training (**Fig. 3D**; effect of time F (8, 198) = 12.61, p<0.0001; effect of genotype F (1, 198) = 5.946, p=0.0156) and displayed greater swim speed (**Fig. 3E**; effect of time F (8, 198) = 7.604, p<0.0001; effect of genotype F (1, 198) = 11.63, p=0.0008).

**Figure 3.**
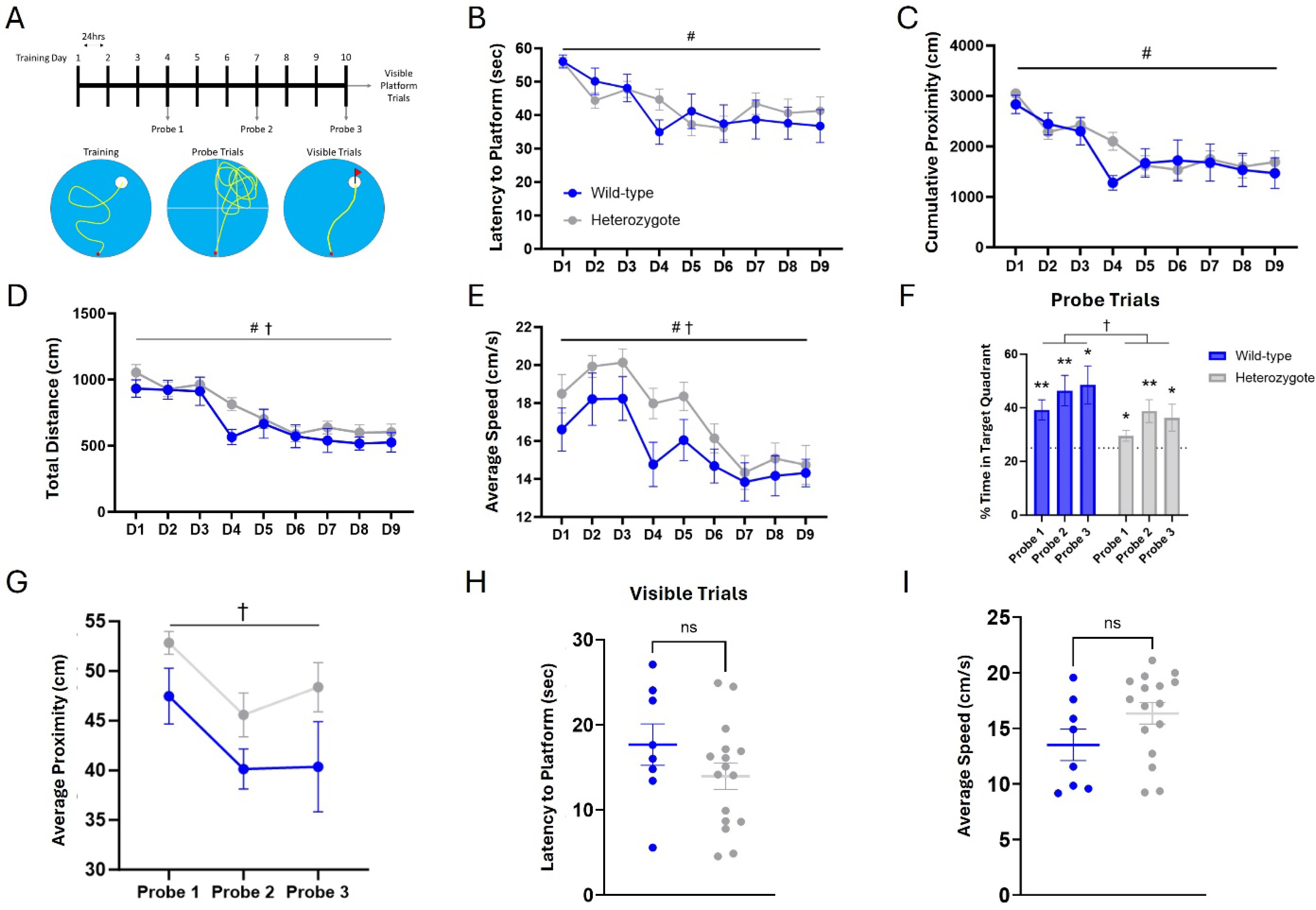
*DSCAM*^2J^+/− heterozygotes display spatial memory deficits in the Morris water maze (MWM) (**A**) Training consisted of nine consecutive days of four trials per day in which mice were allowed to explore for 60s or until they reached the hidden platform, which is set just below the surface of the water. (**B**) During training, the latency to reach the platform significantly decreased across days with no significant differences between groups. (**C**) The cumulative proximity, which is a measure of proximity to the platform location during each trial, also decreased with training days but resulted in no difference between groups. (**D, E**) While there was no difference between the groups in respect to distance travelled during training, the average speed was significantly increased in the heterozygotes. Three probe trials were given to test memory of the platform location. During these trials, the platform was removed from the pool and the mice were given 60s to search for the now removed platform, measuring the percentage of time spent searching in the correct quadrant of the pool as a measure of spatial memory. (**F**) Both groups performed significantly better than chance in all three probe trials, but WT outperformed heterozygotes across probes. (**G**) The average proximity to the correct platform location was significantly increased in heterozygotes. To ensure there were no other behavioral factors present in the groups that might impact performance, on Day 10 the mice received six trials where the platform was highlighted by a visible flag. (**H, I**) Both groups were able to quickly swim to the platform, showing no significant differences in visual recognition or swim speed. Results are presented as Mean ± SEM. Sample size: *DSCAM*^2J^+/− n = 16 and WT n = 8. Statistical tests are as follows: (**B-E**) Two-way ANOVA, where # denotes main effect of training and † denotes main effect of genotype. (**F**) One-sample *t*-test compared to chance = 25% and two-way ANOVA where † marks main effect of genotype. (**G**) Two-way ANOVA, where † denotes main effect of genotype. (**H, I**) Two-tailed *t*-test. Significant differences are shown (* p < 0.05 and ** p < 0.01).

To test their memory recall, the mice were evaluated three times throughout the training procedure in probe trials. Compared against chance (25%), both groups showed a preference for exploring the target quadrant (**Fig. 3F**; wild-type P1 t=3.810, df=7, p=0.0066; wild-type P2 t=3.769, df=7, p=0.007, wild-type P3 t=3.324, df=7, p=0.0127; heterozygote P1 t=2.267, df=15, p=0.0386; heterozygote P2 t=3.252, df=15, p=0.0054; heterozygote P3 t=2.258, df=15, p=0.0393). However, when the groups were compared against each other the heterozygotes display significantly poorer performance (**Fig. 3F**; effect of genotype F (1, 22) = 4.625, p=0.0428). Furthermore, heterozygotes showed an increase in their average proximity to the platform location during the probe trials (**Fig. 3G**; effect of probe F (1.890, 41.59) = 5.191, p=0.0108; effect of genotype F (1, 22) = 6.584, p=0.0176). These results suggest a moderate deficit in spatial learning and memory in the *DSCAM* heterozygotes.

To determine whether there were any visual or locomotor differences that could serve as confounds, on the final day of the experiment the visual platform trials were run. The heterozygotes showed no significant difference in latency to find the platform compared to wild-type controls (**Fig. 3H**; t=1.343, df=22, p=0.1931) nor did they display a significant difference in swim speed (**Fig. 3I**; t=1.690, df=22, p=0.1051). These results demonstrate that the poor performance exhibited by the heterozygotes was not due to visual or locomotor deficits.

### Loss of one copy of *DSCAM* impairs Pavlovian fear learning

Tone-cued and contextual fear conditioning were used to assess Pavlovian fear learning and memory. The 5-day protocol includes three days of tone-shock paired training, contextual recall 24 hours later, tone response testing in a novel context on Day 5, and a final contextual recall at 30 days (**Fig. 4A**). During training days, mice experience a 3-minute baseline period that can be used to establish a learning curve. While both groups did display learning, heterozygotes showed a significant deficit in baseline freezing (**Fig. 4B**; effect of time F (1.306, 27.42) = 163.6, p<0.0001; effect of genotype F (1, 21) = 32.74, p<0.0001; post-hoc comparison of days D1 p>0.9999, D2 p=0.7074, D3 p=0.0010). This freezing deficit was present in the 24-hour context recall test (**Fig. 4C**; t=3.254, df=21, p=0.0038). Thirty days after the final training day, mice were subjected to a context recall test again, where heterozygotes still displayed a deficit in contextual fear recall (**Fig. 4D**; t=2.989, df=21, p=0.007). A comparison of the 24-hour and 30-day context recall showed heterozygotes had a substantial reduction in freezing at the level of the individual subject (**Fig. 4E**), suggesting a lack of long-term consolidation of the fear memory. In tone-cued fear conditioning in a novel context both groups exhibited a freezing response in response to the tone (**Fig. 4F**; effect of test F (1, 21) = 26.15, p<0.0001; wild-type p=0.0192, heterozygote p=0.0002) but the tone responses were not statistically different between groups (**Fig. 4F**; post-hoc comparison of tone test p=0.1798). Both groups showed a significant reduction in freezing behavior when introduced to a novel context (**Fig. 4G**; effect of context F (1, 21) = 24.06, p<0.0001; wild-type p=0.0069, heterozygote p=0.0021) but wild-type mice exhibited a heightened generalization in the novel context (**Fig. 4G**; effect of genotype F (1, 21) = 16.11, p=0.0006; Context B p=0.0104). Overall, these results demonstrate a considerable contextual fear learning deficit in the heterozygotes.

**Figure 4.**
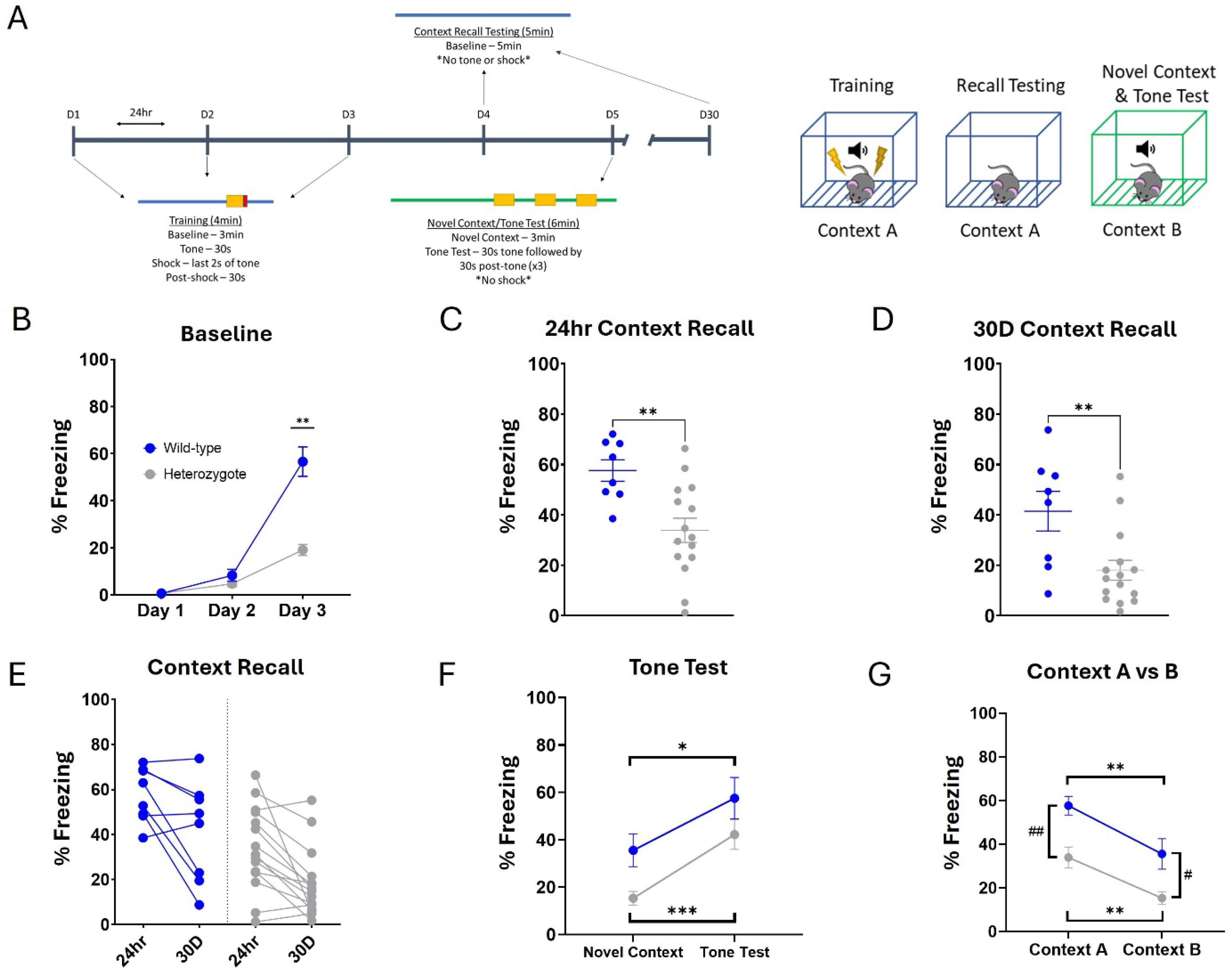
*DSCAM*^2J^+/− heterozygotes exhibit long-lasting deficits in contextual fear learning and memory. (**A**) For three days, mice were placed in a training context (Context A) and allowed to explore for 180s before a 30s tone is played that co-terminated with a foot-shock (2s, 0.75mA). Following training, contextual recall was carried out 24hr later in the same context (Context A). To test tone recall, on Day 5 mice were introduced to a completely novel context (Context B), allowed to explore for 180s after which the 30s tone is repeated 3x with 30s inter-tone intervals. Thirty days after the last training day, mice were reintroduced to the training context (Context A) to be tested for extended contextual recall. (**B**) During training, heterozygotes displayed significant deficits in fear learning, measured by percent freezing, that became evident by the third training day. (**C**) Heterozygotes exhibited significantly less freezing during the contextual recall. (**D, E**) Furthermore, the deficit in fear learning and memory was present 30 days later, as many of the heterozygotes showed substantial decrement in freezing during the 30-day contextual recall. (**F**) Both groups displayed intact tone recall in the novel context/tone test on Day 5. (**G**) Likewise, both groups exhibited a significant reduction in freezing levels in the novel context (Context B). Results are presented as Mean ± SEM. Sample size: *DSCAM*^2J^+/− n = 15 and WT n = 8. Comparison between heterozygotes and WT mice was analyzed by a two-way ANOVA with planned post-hoc comparisons and/or two-tailed *t*-test. (**G**) Two-way ANOVA, where * denotes an effect of test and # denotes an effect of genotype. Significant differences are shown (* and # p < 0.05, ** and ## p < 0.01, **** p = 0.0001).

To examine an alternative form of fear learning that integrates fear-motivated decision-making, mice were evaluated in the passive avoidance task. This task incorporates associative learning and the suppression of the innate desire to enter the dark environment to avoid an aversive stimulus. The 4-day protocol begins with habituation to the chambers followed by two days of acquisition and ends with a retention testing session (**Fig. 5A**). During habituation, mice were given three trials to acclimate to the chambers. Importantly, there was no inherent difference between the groups in their willingness to cross over to the dark chamber once the dividing door opens (**Fig. 5B**; effect of genotype F (1, 21) = 0.6785, p=0.4194). However, there was also no notable difference between the groups in the acquisition and retention testing phases (**Fig. 5C**; effect of genotype F (1, 21) = 0.02969, p=0.8648). These results indicate that the heterozygotes are capable of integrating an instrumental response (i.e. avoidance) with Pavlovian aversive conditioning to the same extent as wild-type mice.

**Figure 5.**
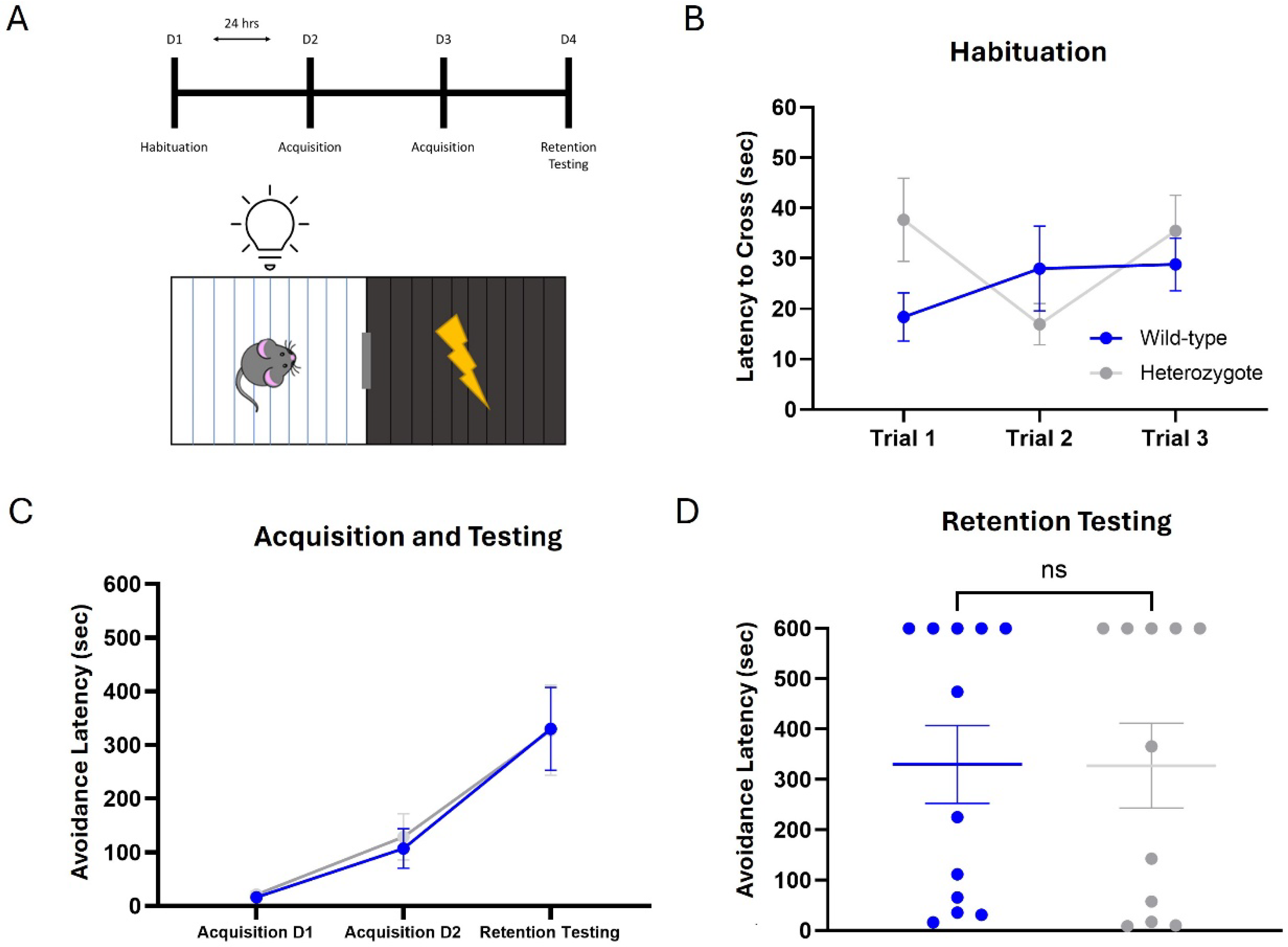
*DSCAM*^2J^+/− heterozygotes do not display deficits in fear-motivated decision-making. (**A**) On the first day, mice were allowed to explore and habituate to both the illuminated and dark chamber. Following habituation, two days of acquisition were completed in which mice were subjected to an unsignaled foot-shock (3s, 0.4 mA) after crossing into the dark chamber. Retention testing was performed 24hrs later. The latency to cross, or the avoidance latency, was recorded as a measure of willingness to enter the dark chamber. (**B**) Both groups were willing to cross into the dark chamber and did so equally across three habituation trials. (**C, D**) During acquisition and retention testing, both groups exhibited an increase in avoidance latency but there were no differences between the groups. Results are presented as Mean ± SEM. Sample size: *DSCAM*^2J^+/− n = 11 and WT n = 12. Comparison between heterozygotes and WT mice was analyzed by a two-way ANOVA with planned post-hoc comparisons and/or two-tailed *t*-test.

### Visual procedural learning is intact in *DSCAM* heterozygotes

The Bussey-Saksida Touch Systems were used to evaluate complex procedural learning as well as visual discrimination. The mice had to first complete the four stages of pretraining (see methods). Pretraining results showed no differences between the genotypes in survival curves when analyzed based on day-by-day performance changes (**Fig. 6A**; Mantel-Cox X^2^(1) = 1.102, p= 0.2939). In a direct comparison of the total number of days required to complete pretraining, there was no difference found between the groups either (**Fig. 6B**; t=1.303, df=25, p=0.2044). However, the heterozygotes did demonstrate a decrease in the number of trials needed to progress through pretraining (**Fig. 6C**; t=2.677, df=25, p=0.0129). This suggests that the heterozygotes may have been more efficient with each training session compared to the wild-type mice, using fewer overall trials to learn each stage.

**Figure 6.**
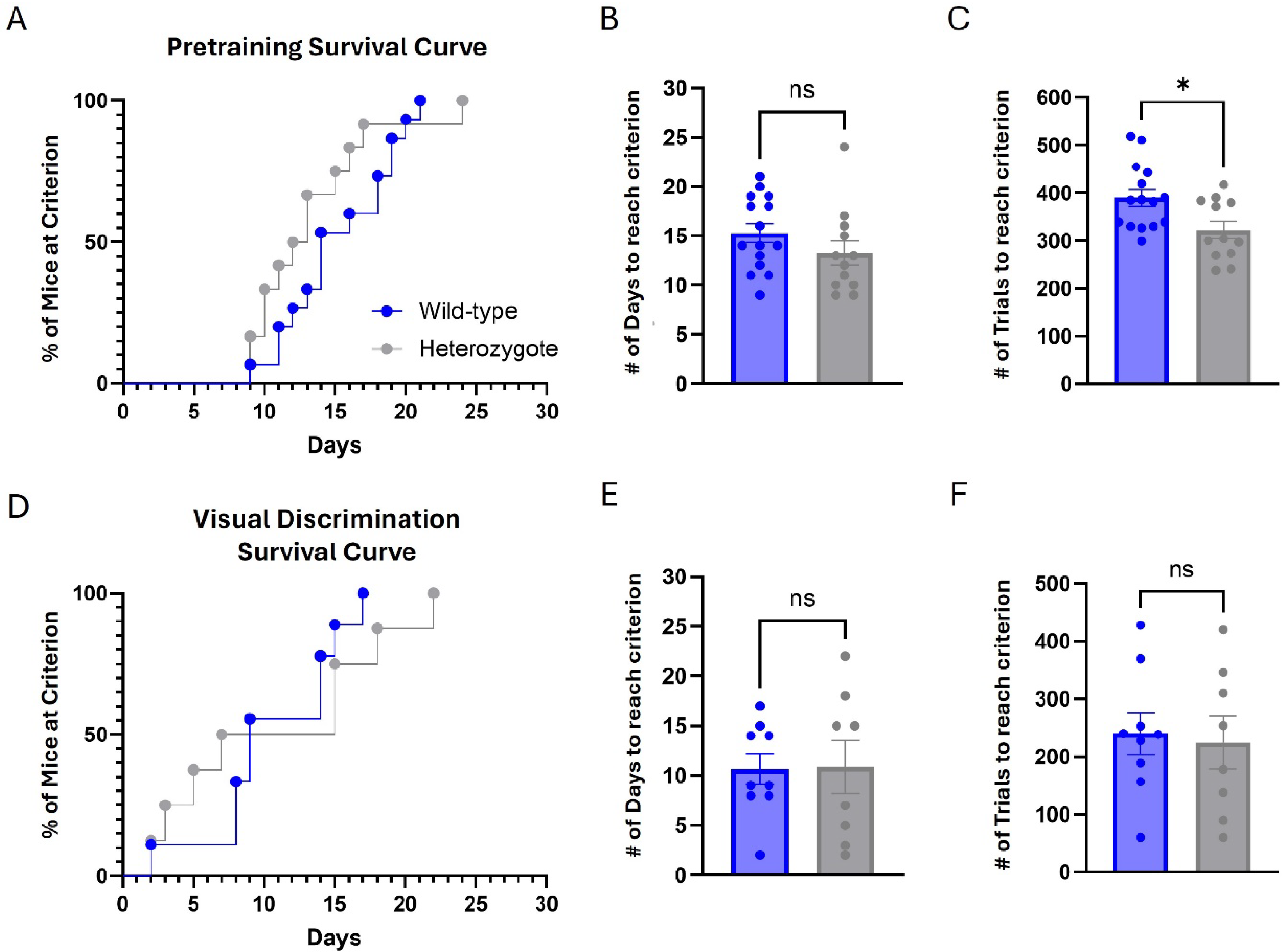
*DSCAM*^2J^+/− heterozygotes exhibit normal procedural learning and higher-level discriminatory learning. Procedural learning and visual discrimination were assessed in Bussey-Saksida Touchscreen systems in daily sessions until criterion was reached. (**A**) Pretraining presented as a survival curve indicated no gross differences between groups in the number of days to reach criterion. (**B, C**) Heterozygotes and WT mice required similar numbers of days to reach criterion during pretraining but needed less trials within those days to progress. Following pretraining, mice were assessed in a pairwise visual discrimination task. (**D**) The survival curve for visual discrimination revealed no significant differences in the number of days to reach criterion, set at 70%. (**E, F**) Similar to pretraining, both groups required nearly the same number of days and trials to complete the task. Results are presented as Mean ± SEM. During pretraining, sample size *DSCAM*^2J^+/− n = 12 and WT n = 15. For visual discrimination, several mice in each group did not complete the task within the given timeframe and were removed from the analysis, leading the sample size to be reduced to *DSCAM*^2J^+/− n = 8 and WT n = 9. Comparison between heterozygotes and WT mice was analyzed by a Log-rank (Mantel-Cox) test (**A and D**) or two-tailed *t*-test.

Once mice satisfied the pretraining stages, they were advanced to the visual discrimination task. In this task, mice were rewarded for interacting with one image (S+) and punished with a timeout period for interacting with a different image (S-). Comparison between the groups revealed no significant difference in performance, as illustrated by a survival curve (**Fig. 6D**; Mantel-Cox X^2^(1) = 0.5957, p=0.4402). Similarly, there was no difference noted in the total number of days to reach criterion (**Fig. 6E**; t=0.06926, df=15, p=0.9457) or in the number of trials (**Fig. 6F**; t=0.2771, df=15, p=0.7855). These results suggest that there was no difference between the wild-type and heterozygotes in the ability to visually discriminate between two distinct images.

## Discussion

In this study, we demonstrate that reduced expression of *DSCAM*, due to the heterozygosity of a LoF mutation, can produce several behaviors that resemble those observed in children and adults with ASD. First, loss of one copy of *DSCAM* led to a significant increase in activity and anxiety-like behavior when exploring a novel environment, and produced difficulties in motor coordination. Additionally, the mutant mice displayed deficits in spatial learning and memory across multiple domains, including short (i.e., working) and long-term memory. Third, fear learning was acutely affected in *DSCAM* heterozygous mice, mirroring the lack of awareness to danger that is reported in children with ASD. However, decreased *DSCAM* expression did not result in a deficit in procedural learning. The close similarity to behaviors observed in ASD, particularly in children, suggests that *DSCAM* LoF mutations and the mechanisms involved may underlie those phenotypes.

### Hyperactivity

Diagnosis of ASD typically occurs in early childhood, often by 3-5 years old (Van ’T Hof et al., 2021). Multiple screening tests exist, but all require a thorough behavioral evaluation which is often repeated at several key stages of development to confirm the diagnosis (Thabtah and Peebles, 2019). However, many of the symptoms display considerable overlap with other commonly diagnosed childhood disorders such as attention-deficit hyperactivity disorder (ADHD), obsessive-compulsive disorder (OCD), and bipolar disorder (BD) (Rosen et al., 2018). It is not uncommon for ASD to be co-diagnosed with one or more of these disorders. In fact, it has been reported that 40-70% of ASD cases qualify and are co-diagnosed for ADHD (Antshel and Russo, 2019). One of the primary reasons for the overlap is hyperactivity, which is distinctly present in ADHD but also a prominent symptom of ASD. Therefore, this is a key phenotype to recapitulate in a mouse model of ASD. Here, we found that that *DSCAM* heterozygous mice travelled significantly more distance during exploration of the novel environment (**Fig. 1A, B**). Hyperactivity was also noted in the Y-maze with increased number of arm entries and distance travelled (**Fig. 2B, C**), and in the Morris water maze with an increased swim speed (**Fig. 3E**). Altogether, these results show that loss of one copy of *DSCAM* leads to hyperactivity in exploratory behavior akin to that observed in childhood ASD.

### Motor Coordination

Individuals with ASD have long been documented as expressing peculiar motor behaviors, specifically restricted repetitive behaviors (RRBs) such as body rocking or hand flapping (Loftin et al., 2008). These RRBs are a core feature of the DSM-V criteria to qualify for ASD diagnosis (Lord and Bishop, 2015). However, the motor phenotype of ASD extends beyond these repetitive movements. There is evidence that broader impairments in both gross and fine motor skills are present in children with ASD (Lloyd et al., 2013). Furthermore, these impairments may correlate with the degree of intellectual disability (Bhat, 2021), with more cognitively impaired individuals displaying more significant alterations in motor skills. The results of these studies, among others (Kaur et al., 2018, Bremer and Cairney, 2018), strengthen the argument that motor performance should be tested and included in the diagnostic criteria for ASD. Therefore, it is important that motor behavior impairments be expressed and studied in mouse models of ASD. We evaluated our *DSCAM* heterozygous mice in the rotarod behavioral tasks to probe their locomotor coordination. The mice had substantial difficulty with the rotarod task and were not able to improve upon their performance with additional training (**Fig. 1E, F**). As this task required active balance and adjustment to an accelerated rotation, any impairment in motor skills or praxis will negatively affect performance. These results align with reported studies that demonstrate motor coordination deficits in children with ASD.

### Short- and Long-term Memory

Despite the common misconception, autism is not often coupled with savant syndrome. Estimates suggest only 1 in 10 individuals with ASD show savant-like traits (Treffert, 2014). In fact, intellectual disability (ID), of varying severity, is more frequently observed in ASD and cooccurring rates may be as high as 50-70% (Matson and Shoemaker, 2009). Cognitive impairments vary amongst individuals, but the majority of ASD cases with ID show deficits in domains such as executive function, language, working memory, and spatial and episodic memory (Banker et al., 2021). This type of impairment profile strongly suggests the prefrontal cortex and hippocampus as brain regions that may be affected in ASD. The first area that we sought to evaluate in the *DSCAM* heterozygous mice was working memory in the Y-maze (**Fig. 2**), where we observed deficits in correct alternations that placed them significantly below chance (**Fig. 2D**). Furthermore, the heterozygotes had more direct revisit errors (**Fig. 2E**), signaling that these mice were unable to form a proper working memory map of the maze – regardless of the spatial cues available to them. It is worth noting that the *DSCAM* heterozygous mice were hyperactive during the task, with a greater number of arm entries and distance travelled (**Fig. 2B, C**). This may have also contributed to their poor performance, if their hyperactivity had impeded their ability to formulate a cognitive map of the maze. It is therefore important to consider that ADHD, with which ASD is often co-diagnosed, also carries substantial impairments in working memory and it may be difficult to separate the two disorders in this domain (Truedsson et al., 2020).

While working memory deficits are well-documented in ASD, deficits in long-term memory, specifically within the realm of spatial and episodic memory, are less understood. Children and adults with ASD typically show difficulties within these domains, such as an inability to retrieve details of life events and often not including themselves in stories from the past. The source of this impairment had been attributed to the social cognition deficits that are a hallmark of ASD (Wantzen et al., 2021, Robinson et al., 2017), but new research suggests that there are distinct disruptions in explicit learning and memory that may be independent of those reported in social cognitive processes (Agron et al., 2023, Ring et al., 2015). If so, focusing on the hippocampus and behaviors specific to disruptions of hippocampal structure and processes may bear fruit for understanding ASD cognitive deficits. In this study, the *DSCAM* heterozygous mice did show a subtle impairment in the Morris water maze task (**Fig. 3**), which is heavily dependent on the hippocampus. While the mutant mice were able to perform the task and significantly outperform chance in the probe trials, they exhibited a less selective search strategy (**Fig. 3F, G**). These results demonstrate that *DSCAM* heterozygotes can still learn and recall the platform location but less effectively so, which is reminiscent of ASD individuals’ ability in episodic memory tests. When coupled with Chen et al.’s findings that a complete knockout of *DSCAM* produces social recognition impairments (Chen et al., 2022), these results agree with the theory that social cognitive and spatial learning deficits coexist in ASD and may be independent phenotypes.

### Fear Learning

Another aspect of learning that appears to be differentially affected in ASD is fear learning, and a common symptom of ASD in children is a “muted” fear response (Macari et al., 2018). This behavior is difficult to interpret as it could be due to multiple factors. There are known impaired domains in ASD that could be playing a role, such as poor social recognition and emotional dysregulation (Macari et al., 2021) or a deficit in spatial learning implicating the hippocampus. There are also less studied aspects of ASD that may be contributing factors, such as improper consolidation of fear memories in the anterior cingulate cortex or a more direct disruption of the emotional learning process within the amygdala. All of these components are involved in fear circuitry (Shin and Liberzon, 2010), and it seems possible that a combination of affected components is likely. Brain imaging studies have shown that adults with ASD exhibit a heightened anxiety and increased reactivity in the amygdala while in safe contexts, and an inability to properly differentiate between safe and threatening conditions (Top Jr et al., 2016). Researchers postulate that this may be due to a saturation of the amygdala activity through heightened baseline arousal, and therefore fear cannot be appropriately contextualized. Moreover, there appears to be an inverse correlation between ASD symptom severity and the strength of the fear response (South et al., 2011), suggesting a direct relationship between the behavioral symptoms and possible disruption of the neurocircuitry in the amygdala-hippocampus-cortex fear network. The molecular underpinning of this relationship needs further investigation, but fear learning does indeed appear to be impaired in ASD. In this study, we detected a significant deficit in fear learning and memory within the *DSCAM* heterozygous mice that extended to thirty days post-exposure (**Fig. 4B-E**). The experimental and control groups displayed similar responses to the cued tone (**Fig. 4F**), signaling that the deficit is likely in context-specific memory, and therefore likely dependent on the hippocampal processes that are encoding those context-based memories. Determining the extent to which the amygdala may play a role in this phenotype, and whether loss of one copy of *DSCAM* drives disruptions in the amygdala among other regions in fear neurocircuitry would be of great interest and could inform mechanisms in ASD.

### Implicit Learning

ASD is a neurodevelopmental disorder that affects the way individuals process the world, both socially and non-socially, to learn and make decisions. The previously discussed studies outline the varying degrees of cognitive impairment present in ASD, impacting numerous domains.

Nonetheless, many autistic individuals succeed in academic settings and beyond, despite these learning difficulties. This may be explained by the fact that implicit memory, such as perceptual and procedural memory, is seen to be unaltered in ASD (Foti et al., 2015, Boucher and Anns, 2018). Combining these findings, a “see-saw effect” hypothesis has emerged in the field, suggesting that intact implicit memory systems act to compensate for the impairment seen in explicit systems such as episodic memory (Boucher and Anns, 2018). With ASD rising in the general population (Maenner et al., 2023) there is an increased demand to understand the disorder and develop strategies and methods to accommodate ASD children in school (Paisley et al., 2023) and adults in the workplace (Johnson et al., 2020). An example of which may be a greater focus on active learning and allowing autistic individuals to learn at their own pace. Indeed, active learning strategies have been seen to improve episodic memory in children with ASD (Fantasia et al., 2020). This may help explain our findings in the touchscreen systems (**Fig. 6**), where the *DSCAM* heterozygous mice performed as well as, or even slightly better than, the control mice. With procedural learning intact, the pretraining stage was not an impediment and the *DSCAM* heterozygotes even completed this stage in less overall trials. While not every mouse completed the visual discrimination portion of the experiment, there was no difference between each group’s performance in those that did complete the experiment (**Fig. 6D-F**). The nature of this task, where each daily session allows the mouse to be undisturbed for an hour and learn at their own pace, may have provided the right conditions for optimal performance despite previously illustrated cognitive deficits, as has been argued for human cases of ASD. Alternatively, it is possible that the trend suggesting that the *DSCAM* heterozygous mice outperformed their wild-type littermates reflects some sort of perseverative tendencies in the heterozygotes.

### Conclusions

Growing evidence shows that *DSCAM* acts in a dose-dependent manner. While the overexpression of *DSCAM* may be contributing to cognitive impairments observed in Down syndrome, its underexpression could be a pathway to autistic-like phenotypes. Our results established that *DSCAM* heterozygous mice exhibit many behaviors that are reminiscent of those observed in ASD, such as hyperactivity, motor coordination abnormalities, and learning and memory deficits. These findings, coupled with previous research, provide compelling evidence that *DSCAM* LoF mutations may be an avenue through which ASD-related behaviors are produced and advocate for *DSCAM*^2J^+/− as a model for ASD research and therapeutic testing.

## Materials and Methods

### Animals, breeding, and genotyping

For all experiments, littermates were age-matched and divided into cohorts (**Table 1**). The *DSCAM*^2J^ (C3H/HeDiSn-*DSCAM^2J^*/GrsrJ) mice were originally acquired from Jackson Laboratories (Strain #: 006038) and migrated to a C3H background by crossing *DSCAM*^2J^+/− mice to sighted C3H (C3Sn.BLiA-*Pde6b*^+^/DnJ), also acquired from Jackson Laboratories (Strain #: 003648) for 10+ generations. *DSCAM*^2J^ were then maintained by the same cross to produce heterozygous (+/−) and wild-type (+/+) littermates. Therefore, *DSCAM*^2J^+/− will also be referred to as heterozygotes and *DSCAM*^2J^+/+ as wild-type. Mice were housed by sex, at a maximum of 5 animals per cage, in a temperature-controlled vivarium (22°C) with a 14/10-hour light/dark cycle until they had reached the designated age range to begin studies. Mice were provided *ad libitum* food and water, except during visual discrimination experiments as detailed. All mice were 2-6 months of age during the experiments and equal numbers of males and females were used.

All genotyping was performed on tail tips by Transnetyx (Memphis, TN) based on protocols developed by Jackson Laboratories. All mouse work and protocols were approved by the University of Michigan Institutional Animal Care and Use Committee (IACUC) and in accordance with the National Research Council Guide for the Care and Use of Laboratory Animals (NIH).

### Behavioral tests

All experiments were conducted with the investigator blind to genotype. Cohort sample sizes were determined based on need of the subsequent experiments, and estimates were calculated with a detection level of ∼10% using G*Power software (Faul et al., 2007) with α set to 0.05 and the power (1-β) value set to 0.95. Standard deviation values were derived from historical lab data on the included behavioral tasks.

#### Open-field exploration

To assess exploratory behavior, including activity levels, open-field experiments were conducted. Experiments were carried out in a rectangular open arena (74 x 74 x 29 cm) with smooth white opaque walls and floor made of acrylic. The arena was illuminated at approximately 45 lumens, measured at the center. Each mouse was individually placed into the center of the arena and allowed to freely explore the environment for 10 minutes, after which the mouse was removed and returned to their home cage. Between each mouse the arena was cleaned with 70% ethanol. LimeLight 3 video tracking software (Actimetrics, Evanston, IL) and a camera mounted above the arena were used to record behavior during the trial. Total distance traveled, time spent in the inner zone of the arena (46 x 46 cm), and the number of crossings between zones was analyzed offline. One mouse was removed due to inactivity that was statistically determined to be an outlier.

#### Rotarod

Locomotor coordination and activity were evaluated using the rotarod (TSE Systems, Chesterfield, MO). Mice were placed on the rotating rod, initially moving at a constant rate of 2 rpm. Following a 10s habituation period to allow the mice to get accustomed to initial rotation, the rod was then accelerated from 4 to 40 rpm at a constant rate. The maximum trial duration was set to 5 minutes. Mice were run one at a time and given a single trial each day for three consecutive days. Latency to fall was recorded by the TSE RotaRod software using the pressure plate in the flooring of the apparatus and manually by the experimenter as a backup.

#### Hot plate

Sensory response to thermal nociception, i.e., pain sensitivity, was measured using the hot plate test (IITC Life Sciences, Woodland Hills, CA). The black aluminum surface (27 x 30 cm) was maintained at 50 ± 0.5°C and confirmed to be stable using a thermistor probe connected to the hot plate apparatus. An acrylic cylindrical enclosure (15 cm height, 10 cm inner diameter) was used to limit the movement of the mice on the heated surface during the experiment. Each mouse was placed onto the surface and allowed to explore while being closely monitored by an observer. The latency to hindlimb lick was determined and recorded as the response time, at which point the mouse was promptly removed from the enclosure and returned to their home cage.

#### Y-maze

Working memory was measured by spontaneous alternation in the Y-maze. The Y-maze was built with three symmetrical arms (7.5 x 18 x 13 cm) at a 120° angle. The maze was constructed of rigid white opaque acrylic. Connecting the three arms is an equilateral triangular space (7.5 cm). Each trial began with the mouse being placed in the center triangular space facing the same arm. The mouse was then allowed to explore the maze for 8 minutes. An arm entry was recorded when the center-point of the mouse crossed the half-way point of the arm. Triads were recorded, which is defined as a set of three arm entries. A correct triad was determined to be consecutive entries into the three different arms of the maze, and any other combination was considered an error. Direct revisits were recorded when a mouse leaves an arm, enters the center zone, and returns to the same arm again. Indirect revisits were defined as when a mouse leaves an arm, passes through the center zone, enters another arm, and then returns to the original arm. Behavior was recorded using an overhead camera and analyzed through EthoVision XT v17 software (Noldus, Wageningen, the Netherlands).

#### Morris water maze

Long-term spatial learning and memory was assessed using the Morris water maze (MWM). The arena was a 1.2-meter diameter pool filled with water that was semi-opaque through the addition of white nontoxic paint. The water temperature was maintained at 25±2 C° using an electric heating mat underneath the pool. For each of the training trials, a circular platform (10 cm diameter) was placed in the NE quadrant of the pool and sat approximately 1 cm below the water surface. Each training session began with the mouse being placed on the platform for ∼10s before starting their first trial. The mouse was then placed into the water along the wall and allowed to search the arena for the platform. The training trial ended when the mouse reached the platform or when 60s had elapsed. Mice were trained for four trials per day for 9 consecutive days, and the starting position for each trial was chosen pseudo-randomly from seven potential positions. On training days 4, 7, and 10, the mice were evaluated in a probe trial that was conducted before any training trials on the day. The probe trial removed the platform from the pool to assess their memory of the platform location. Mice were placed opposite the platform’s usual location and given a 60s trial to search the arena. On the last day of the experiment and following the final probe trial, visible platform trials were conducted to assess locomotor and visual capabilities. The visible trials consisted of six trials where the platform now had a visual cue (a paper flag) to mark the platform location. Trials were run in sets of 2, where the platform was changed to a different quadrant for each set. Training trial duration was averaged together for each day to represent that mouse’s performance, where applicable. Behavior was recorded by an overhead camera at 15 fps and analyzed through the WaterMaze software (Actimetrics, Evanston, IL).

#### Fear conditioning

Contextual and cue-based fear conditioning was conducted to assess long-term fear learning and memory. Four operant chambers (Med Associates, Fairfax, VT) were organized on a steel rack in an isolated room. The chamber was constructed of clear acrylic on the front, back, and top, and aluminum on the two remaining sides. The floor was made of stainless-steel parallel bars forming a grid with a metal tray positioned below. Each of the metal floor grids was connected to a shock controller, which was managed by a desktop computer running FreezeFrame software (Actimetrics, Evanston, IL). A camera was positioned above each chamber to record mouse behavior back to the same computer. The primary measure was freezing, which was assessed using the FreezeFrame software to compare the movement of the mouse frame by frame.

The surrounding environment was altered depending on the chosen context. Two contexts were used throughout the experiments, designated Context A and Context B. Context A consisted of white room lighting, white noise from a sound machine, 70% ethanol as a cleanser and an odor, and a floral-patterned curtain that covers the chambers during the session. Context A was used for the training and context recall components of the experiment. Context B was significantly modified to act as a novel environment in comparison to Context A. Therefore, Context B used red room lighting, no added noise, 2% acetic acid as a cleanser and odor, and a blue and white striped curtain. Additionally, the metal floor grid was covered with a white acrylic sheet and a cushioned mat flooring, and a curved sheet of white acrylic was inserted into the chamber to alter its dimensions.

Using the four chambers, mice were transferred from their home cages to the chambers individually. The experiment was conducted on consecutive days, with the first three days acting as conditioning training. For conditioning sessions, mice were placed into Context A for 3 minutes prior to a 30s tone (2.8 kHz, 75 dB) whose termination aligned with a foot-shock (2s, 0.75 mA). Thirty seconds later, the mice were removed from the chamber. To test contextual memory, 24 hours after the final conditioning session the mice were returned to Context A for a 5-minute period in the absence of any tone or foot-shock. Finally, to test cued memory, 24 hours after the context test the mice were placed in Context B for 3 minutes, after which the 30s tone was initiated three times with a 30s inter-tone interval. Mice were then returned to their home cage.

During training, one mouse escaped from the chamber and was pursued by the investigator until caught. This mouse was subsequently removed from the analysis due to a distinct experience that could be fear inducing, which follows standard exclusion criteria for this behavioral task.

#### Passive avoidance

Fear learning, in which the animal must choose between approach or avoidance, was evaluated using the Passive Avoidance paradigm. The apparatus (Maze Engineers, Shokie, IL) consisted of two chambers (20 x 20 x 20 cm) that were connected via an automated sliding door (7 x 5 cm). The chambers were constructed of black acrylic walls, a clear acrylic lid, and stainless-steel grid floor. One chamber was well-lit, with the lid being kept clear, and a bright light was present within the chamber to invoke photophobia. The second chamber was kept dark, in which the lid was shrouded in black construction paper and the light was not turned on. During the habituation phase, the mouse was introduced to the light chamber and allowed to explore for one minute, at which point the sliding door was opened to reveal access to the dark chamber. The latency to cross was recorded via infrared beams, and the door was closed once the mouse crossed over to the dark chamber. After an additional minute of exploration of the dark chamber, the lid was removed, and the mouse was returned to their home cage. The maximum time to cross was set to 10 minutes. Each mouse was given three such habituation trials on the first day of the experiment. On the second day, referred to as the acquisition trial, the mouse is again allowed to cross to the dark chamber. After one minute an aversive stimulus, a mild foot-shock (3s, 0.4 mA), was delivered through the floor grid and the mouse was removed from the chamber 30s after the foot-shock. Twenty-four hours after the foot-shock was delivered, the mouse was placed into the apparatus again, and the latency to cross to the dark chamber was recorded in the Passive Avoidance software. A longer latency is correlated with a stronger association of the dark chamber with the aversive stimulus.

#### Touchscreen and visual discrimination

Procedural learning and higher-level discriminatory learning was assessed using the Bussey-Saksida Touchscreen system (Lafayette Instrument, Lafayette, IN). To avoid neophobia, the liquid reward, strawberry Ensure™ milkshake, was introduced to the home cage *ad libitum* for three days prior to the experiment beginning. Before the mice could be tested in visual discrimination, they were taken through a protocol to become adjusted to the touchscreens. However, to provide additional motivation, the mice were food deprived to 90-95% of their body weight. The mice were then accustomed to touchscreen chamber environment for the first time in a ten-minute session. Habituation continued for three additional days, with autotraining to the strawberry milkshake reward (20 µL) in 20 and then two 40-minute sessions. To begin the pretraining phase, mice were then trained to Initial Touch, where images were randomly displayed on the screen and an interaction with the screen elicited 3x the reward (60 µL) and initiated a 10s inter-trial interval (ITI). The Initial Touch phase lasted three days to encourage association between screen interaction and reward. Each mouse then began the Must Touch phase, where interaction with a displayed image was required to elicit a reward, while interaction with a blank screen incurred no penalty. Completing 30 trials within a one-hour period satisfied this phase and advanced the mouse to the Must Initiate phase. Within this phase, mice must first nose-poke to initiate a trial and bring an image onto the screen and then interact with that image for a reward. Successful completion of 30 trials within one-hour progressed the mice to the Punish Incorrect phase, which introduced a 20s timeout penalty for interaction with a blank image location. This phase required a criteria of 24/30 correct trials, or 80%, to complete the pretraining and advance to visual discrimination. All mice progressed through the phases depending on their individual metrics, and therefore their days and trials to criterion/completion were recorded as a measure of their performance. Two of the mice did not complete the pretraining steps, remaining sedentary during the majority of the sessions, and were therefore removed from the study.

Once pretraining was complete, mice began 2-choice visual discrimination. Two images were displayed at once, interaction with the one image was rewarded (S+) and reinforced with strawberry milkshake reward while the other (S-) was punished with a 20s timeout. Interaction with the incorrect image instituted correction trials, where the trial was repeated until the S+ was chosen successfully. Training was continued on weekdays until criterion was reached, which was set at 21/30 correct trials, or 70%, and maintained for 2 consecutive days. Mice were limited to a total of 40 days to complete the visual discrimination task, and any mice that did not complete the task within that timeframe were removed from the analysis. Therefore, 6 wild-type and 4 heterozygotes were removed.

### Statistical analysis

All experiments and analyses were done with the experimenter blind to genotype. Analysis of the data was performed using GraphPad Prism 10. Average data is presented in the figures as the mean ± the standard error of the mean (SEM). The appropriate statistical tests were chosen depending on the experiment and are indicated as such within the figure legends. These tests include: Two-tailed unpaired Student’s *t*-test (for comparison of groups); Mixed-effects model (REML); Log-rank (Mantel-Cox) test; Two-way ANOVA + planned post-hoc comparisons with Bonferroni correction and/or repeated measures (for multifactor comparisons and repeated timepoints); and One-tailed Student’s *t*-tests (for comparison against chance). Statistical significance was considered as p < 0.05.

## Acknowledgements

We thank Drs. Shannon Moore, Yujia Hu, and Sarah Elzinga for helpful discussions and feedback as well as technical support. This work was supported by grants from NIH (R01AG052934 to G.G.M., and R01EB028159 and R21NS094091 to B.Y.), a seed grant from Brain Research Foundation to B.Y., and Protein Folding Disease Initiative of the University of Michigan to G.G.M. and B.Y., and the University of Michigan Rackham Research Grant to R.C.N.

## Author Contributions

R.C.N., B.Y., and G.G.M. conceived of the project and designed the experiments. K.A.S. and T.H. performed the mouse breeding and colony maintenance. R.C.N., K.A.S., and U.B. performed the behavioral experiments and data analysis. R.C.N. wrote the manuscript with B.Y. and G.G.M. providing meaningful edits and comments. Subsequent revisions were made by R.C.N., K.A.S., T.H., B.Y, and G.G.M. Project supervision was done by G.G.M.

## Data Availability

All relevant data can be found within the article and its supplementary information and is available upon reasonable request.

